# Characterization of human-iPSCs derived spinal motor neurons by single-cell RNA sequencing

**DOI:** 10.1101/2019.12.28.889972

**Authors:** Louise Thiry, Regan Hamel, Stefano Pluchino, Thomas Durcan, Stefano Stifani

## Abstract

Human induced pluripotent stem cells (iPSCs) offer the opportunity to generate specific cell types from healthy and diseased individuals, allowing the study of mechanisms of early human development, modelling a variety of human diseases, and facilitating the development of new therapeutics. Human iPSC-based applications are often limited by the variability among iPSC lines originating from a single donor, as well as the heterogeneity among specific cell types that can be derived from iPSCs. The ability to deeply phenotype different iPSC-derived cell types is therefore of primary importance to the successful and informative application of this technology. Here we describe a combination of motor neuron (MN) derivation and single-cell RNA sequencing approaches to generate and characterize specific MN subtypes obtained from human iPSCs. Our studies provide evidence for rapid and robust generation of MN progenitor cells that can give rise to a heterogenous population of brainstem and spinal cord MNs. Approximately 58% of human iPSC-derived MNs display molecular characteristics of lateral motor column MNs, ∼19% of induced MNs resemble hypaxial motor column MNs, while ∼6% of induced MNs have features of medial motor column MNs. The present study has the potential to improve our understanding of iPSC-derived MN subtype function and dysfunction, possibly leading to improved iPSC-based applications for the study of human MN biology and diseases.

## Introduction

Human induced pluripotent stem cells (h-iPSCs) have emerged as a powerful model system to investigate the mechanisms of cell specification, function and dysfunction (reviewed in Haston and Finkbeiner, 2015; Ardhanareeswaran et al., 2017; Amin et al., 2019). Moreover, they offer potential as a physiologically-relevant experimental system for disease modelling and discovery of new therapeutics. A key requirement for fulfilling the full potential of h-iPSCs is the ability to reliably differentiate specific cell types with defined phenotypic traits. It is often the case that functionally specialized subtypes of cells hold the most promise for fundamental biological studies, disease modelling or drug discover efforts. Thus, it is of the utmost importance to establish protocols for the differentiation and deep phenotyping of h-iPSC-derived cells that may represent only a small fraction of cells *in vivo*.

Spinal motor neurons (MNs) are a highly specialized group of neurons that reside in the spinal cord and project axons to muscle fibers to control their contraction. The precise coordination of muscle contractions is absolutely essential to generate complex motor behaviors such as walking, grasping or breathing. To ensure such refined coordination, spinal MNs must acquire specific identities matching the muscle types they will innervate. During MN development, all spinal MNs emerge from progenitor cells in the MN progenitor (pMN) domain. The pMN domain is located in the medial portion of the ventral neural tube, just ventral to the progenitor domain 2 (p2) and dorsal to the p3 domain, which give rise to V2 and V3 interneurons (INs), respectively (Alaynick et al., 2011; Stifani, 2014). After the formation of distinct progenitor domains, pMN progenitors initially acquire a general MN fate in response to the expression of specific transcription factors including PAIRED BOX 6 (PAX6) and OLIGODENDROCYTE TRANSCRIPTION FACTOR 2 (OLIG2) (Ericson et al., 1997; Novitch et al., 2001; Vallstedt et al., 2001). As the spinal cord develops, inductive signals along the rostro-caudal axis further differentiate developing MNs, leading to the formation of anatomically defined motor columns, each containing phenotypically distinct MNs with specific axonal projections to different muscle targets (Prasad and Hollyday, 1991; Tsuchida et al., 1994; Jessell, 2000; Alaynick et al., 2011; Francius and Clotman, 2014; Stifani, 2014).

Loss of MN function is associated with several devastating neurological diseases including spinal muscular atrophy (SMA) and amyotrophic lateral sclerosis (ALS) (Kanning et al., 2010; Nijssen et al., 2017), and it is also a pathological feature of a number of spinal cord injury cases (Alizadeh et al., 2019). Patient-derived iPSCs have been generated from ALS and SMA patients (Ebert et al., 2009; Egawa et al., 2012; Chen et al., 2014; Kiskinis et al., 2014) and a number of differentiation protocols have been developed to obtain patient-derived MNs for disease modelling (*e*.*g*., Li et al., 2005; Qu et al., 2014; Amoroso et al., 2013; Du et al., 2015). Although some of these protocols facilitate the generation of highly pure MN populations (Du et al., 2014), the molecular characterization of these cells is often limited to the use of pan-MN markers through immunostaining approaches (Amoroso et al., 2013; Chen et al., 2014; Maury et al., 2014; Du et al., 2015). Given the heterogeneity of spinal MNs, our limited understanding of the biological properties of h-iPSC-derived MNs, and the possibility that not all types of h-MN subtypes may provide disease-relevant experimental models, there is a critical need to improve our understanding of h-iPSC-derived MN biology and function. Here we describe the results of studies applying single-cell RNA sequencing (sc-RNAseq) to characterize the different cell subtypes obtained after subjecting h-iPSCs to a MN differentiation protocol and we describe the implications of these findings for MN disease modeling and drug discovery efforts.

## Materials and Methods

### Generation of motor neurons from human iPSCs

Human iPSC line NCRM-1 (male) was obtained from the National Institutes of Health Stem Cell Resource (Bethesda, MD, USA). Cells at low passage number were cultured in mTeSR medium (STEMCELL Technologies; Vancouver, BC, Canada; Cat. No. 85850) in 6-cm culture dishes (Thermo-Fisher Scientific; Waltham, MA, USA; Cat. No.130181) coated with Matrigel (Thermo-Fisher Scientific; Cat. No. 08-774-552) until they reached 70%-80% confluence. To generate neural progenitor cells (NPCs), iPSCs were dissociated with Gentle Cell Dissociation Reagent (STEMCELL Technologies; Cat. No. 07174) and split 1:5 on T25 flasks (Thermo-Fisher Scientific; Cat. No. 12-556-009) coated with poly-L-ornithine (PLO; Sigma-Aldrich; Oakville, ON, Canada; Cat. No. P3655) and laminin (Sigma-Aldrich; Cat. No. L2020). The mTeSR medium was replaced with a chemically defined neural medium including DMEM/F12 supplemented with GlutaMax (1/1; Thermo-Fisher Scientific; Cat. No. 10565-018) and Neurobasal medium (1/1; Thermo-Fisher Scientific; Cat. No. 21103-049), N2 (0.5X; Thermo-Fisher Scientific; Cat. No. 17504-044), B27 (0.5X; Thermo-Fisher Scientific; Cat. No. 17502-048), ascorbic acid (100 μM; Sigma-Aldrich; Cat. No. A5960), L-Glutamax (1X; Thermo-Fisher Scientific; Cat. No. 35050-061), and antibiotic-antimycotic (1X; Thermo-Fisher Scientific; Cat. No. 15240-062). This medium was supplemented with 3 μM CHIR99021 (STEMCELL Technologies; Cat. No. 72054), 2 μM DMH1 (Sigma-Aldrich; Cat. No. D8946) and 2 μM SB431542 (Tocris Bioscience; Bristol, UK; Cat. No. 1614). The culture medium was changed every other day for 6 days and the resulting NPCs were then differentiated into motor neuron progenitor cells (MNPCs), and subsequently MNs, as described previously (Du et al., 2015) with minor modifications. Briefly, on day 6 NPCs were dissociated with Gentle Cell Dissociation Reagent and split 1:5 with the same medium described above, supplemented with retinoic acid (RA) (0.1 μM; Sigma-Aldrich; Cat. No. R2625) and purmorphamine (0.5 μM; Sigma-Aldrich; Cat. No. SML-0868) in combination with 1 μM CHIR99021, 2 μM DMH1 and 2 μM SB431542 reagents. The culture medium was changed every other day for 6 days and the resulting MNPCs were characterized by immunocytochemistry. MNPCs were expanded for 6 days with the same medium containing 3 μM CHIR99021, 2 μM DMH1, 2 μM SB431542, 0.1 μM RA, 0.5 μM purmorphamine and 500 μM valproic acid (VPA; Sigma-Aldrich; Cat. No. P4543). To generate MNs, MNPCs were dissociated and split 1:2 with the same neural medium supplemented with 0.5 μM RA and 0.1 μM purmorphamine. Culture medium was replaced every other day for 6 days and the resulting MNs were characterized by immunocytochemistry. For sc-RNAseq, MNs were dissociated, plated on Matrigel-coated T25 flasks, and cultured with the same neural medium supplemented with 0.5 μM RA, 0.1 μM purmorphamine, 0.1 μM Compound E (Calbiochem; Cat. No. 565790), insulin-like growth factor 1 (10 ng/mL; R&D Systems; Minneapolis, MN; Cat. No. 291-G1-200), brain-derived neurotrophic factor (10 ng/mL; Thermo-Fisher Scientific; Cat. No. PHC7074) and ciliary neurotrophic factor (10 ng/mL; R&D Systems; Cat. No. 257-NT-050) for 6 days before single cell suspension preparation.

### Characterization of human iPSC-derived cells by immunocytochemistry

Induced MNPCs and MNs were analyzed by immunocytochemistry, which was performed as described previously (Methot et al., 2018). The following primary antibodies were used: mouse anti-OLIG2 (1/100; Millipore Corp.; Billerica, MA, USA; Cat. No. MABN50); rabbit anti-PAX6 (1/500; Covance; Emeryville, CA, USA; Cat. No. PRB-278P); rabbit anti-HOMEOBOX C4 (HOXC4) (1/150; kindly provided by Dr. Jeremy Dasen, New York University School of Medicine); goat anti-SRY-BOX 1 (SOX1) (1/500; R&D Systems; Cat. No. AF3369; mouse anti-NK2 HOMEOBOX 2 (NKX2.2) (1/100; DSHB; Cat. No. 74.5A5-c); rabbit anti-HOMEOBOX PROTEIN CHX10 (CHX10) (1/10,000; kindly provided by the late Dr. Thomas Jessell, Columbia University); mouse anti-HOMEOBOX PROTEIN HB9 (HB9) (1/30; DSHB; Cat. No. 81.5C10-c); mouse anti-ISLET1 (ISL1) (1/30; DSHB; Cat. No. 39.4D5-c); rabbit anti-LIMB HOMEOBOX CONTAINING 3 (LHX3) (1/100; Abcam; Toronto, ON, Canada; Cat. No. ab14555), goat anti-FORKHEAD BOX PROTEIN 1 (FOXP1) (1/100; R&D Systems; Cat. No. AF4534), and guinea pig anti-SCIP (1/16,000; kindly provided by Dr. Jeremy Dasen). Secondary antibodies against primary reagents raised in various species were conjugated to Alexa Fluor 488, Alexa Fluor 555, or Alexa Fluor 647 (1/1,000, Invitrogen; Burlington, ON, Canada). For quantification, images were acquired using an Axio Observer Z1 microscope connected to an AxioCam camera and using ZEN software (Zeiss). For each culture and each time point (NPCs, MNPCs, MNs), images of >500 cells in 3 random fields were taken with a 20X objective and analyzed with Image J.

### Preparation of induced motor neuron culture single-cell suspensions

Single-cells in suspension were prepared from MNs cultured for 28 days (Day 0 defined as start of NPC induction) as follows. MNs were first washed with 2 mL of dPBS (calcium and magnesium free phosphate buffered saline) containing 0.04% BSA (Sigma-Aldrich; Cat. No. A7906) and dissociated with 2 mL of dissociation reagent containing papain (50U; Sigma-Aldrich; Cat. No. P4762) and Accutase (Thermo-Fischer Scientific; Cat. No. A11105-01). Cells in dissociation reagent were incubated for 15-20 min at 37°C to ensure dissociation of cell clusters before adding 5 mL of DMEM/F12 containing 0.04% BSA and 10% ROCK-inhibitor (1/1000; Sigma-Aldrich; Cat. No. 1254), followed by resuspension by gentle pipetting. Cells in suspension were transferred into a 15 mL falcon tube and centrifuged at 1,300 rpm for 3 min at room temperature. The cell pellet was resuspended in 2 mL of dPBS containing 0.04% BSA and ROCK-inhibitor (1/1000). Cells were centrifuged again at 1,300 rpm for 3 min at room temperature and the cell pellet was resuspended in 500 μL of the above resuspension buffer. A 30 μm strainer was used to remove cell debris and clumps and the single-cell suspension was transferred into a 2 mL Eppendorf tube placed on ice. Cells were counted to evaluate cell concentration and viability before sc-RNAseq. The targeted cell concentration was 700-1,200 cells/μL, as recommended in the 10x Genomics guidelines (https://www.10xgenomics.com/solutions/single-cell/).

### Single cell RNA sequencing and in silico analysis

Cells were sequenced at a single-cell level using the microdroplet-based platform, 10x Genomics Chromium Single Cell 3’ Solution (10x Genomics; Pleasanton, Ca), followed by sequencing on a HiSeq4000 system (Illumina; San Diego, CA) at the McGill and Genome Quebec Innovation Center (https://cesgq.com/en-services). The 10x cDNA libraries were sequenced at a depth of 50,000 reads per cell. The raw sc-RNAseq data (FASTQ files) were first processed using the Cell Ranger pipeline (10x Genomics) to demultiplex and align the sequences to the human reference genome, GRCh38. FASTQ files were aligned and empty droplet 10x barcodes were filtered out via Cell Ranger. Post-Cell Ranger, gene/cell matrices from 5,900 single cells were imported into R. We performed quality control using the Bioconductor-package, scater (McCarthy et al., 2017) to filter out damaged cells based on higher than expected (>3 median absolute deviations) proportions of mitochondrial to nuclear genes and lower than expected (<3 median absolute deviations) genes per cell and transcripts per cell, all of which would suggest that the sequenced cells were dead or damaged and thus leaking nuclear transcripts. The normalization was performed by first clustering the cells using the Bioconductor-package, scran (Lun et al., 2016) before computing cluster-based size factors (scran) to finally normalize the dataset. Next, technical noise in the dataset was removed by first modelling the noise (scran) and then performing principal-component analysis (scran). Later principle components account for very little of the variability in the dataset and are thus assumed to represent the random, technical noise while the endogenous variation of co-regulated genes is assumed to be represented in the earlier principle components. Thus, by removing the later principle components, the dataset can be denoised while preserving the biologically-driven variability of gene expression. All software packages are publicly available at the Bioconductor project (http://bioconductor.org). The workflow used for the analysis was adapted from a published workflow developed to accommodate droplet-based systems such as 10X Chromium (Lun et al., 2016). This workflow will be available on https://github.com/regan-hamel/h-iPSCs-MNI.

## Results

### Generation of an enriched population of human iPSC-derived motor neurons

Spinal MNPCs were generated from the h-iPSCs NCRM-1 line by mimicking *in vitro* sequential steps of MN developmental *in vivo*, including neural induction, caudalization and ventralization of NPCs (Li et al., 2005; Wichterle et al., 2002). Spinal NPCs were induced using a small-molecule cocktail composed of the CHIR99021, DMH1 and SB431542 reagents, as previously described (Du et al., 2015). After 6 days from the start of the induction protocol, 81±4% of the induced cells expressed typical NPC markers, including SOX1, and 70±5% expressed the cervical spinal cord marker HOXC4, indicative of a rostral spinal cell fate (Burke et al., 1995) (Figure 1A and B). NPCs were subsequently ventralized through the addition of the sonic hedhehog (SHH) agonist purmorphamine at a concentration of 0.5 μM, in the presence of 0.1 μM RA, resulting in the generation of MNPCs, with 74±6% of induced cells being PAX6-positive^+^and 76±4% expressing the MNPC marker OLIG2 (Briscoe et al., 2000; Jessell, 2000; Alaynick et al., 2011) at day 12 (Figure 1C and D). As expected, a small proportion of spinal ventral IN progenitors was detected in our cultures, with 1.8±0.3% of the total of differentiated cells expressing the p3 domain marker NKX2.2 (Novitch et al., 2001; Sugimori et al., 2007) and 2.5±0.8% expressing the p2 domain marker CHX10 (Debrulle et al., 2019; Alaynick et al., 2011) (Figure 1E and F).

**Figure 1.**
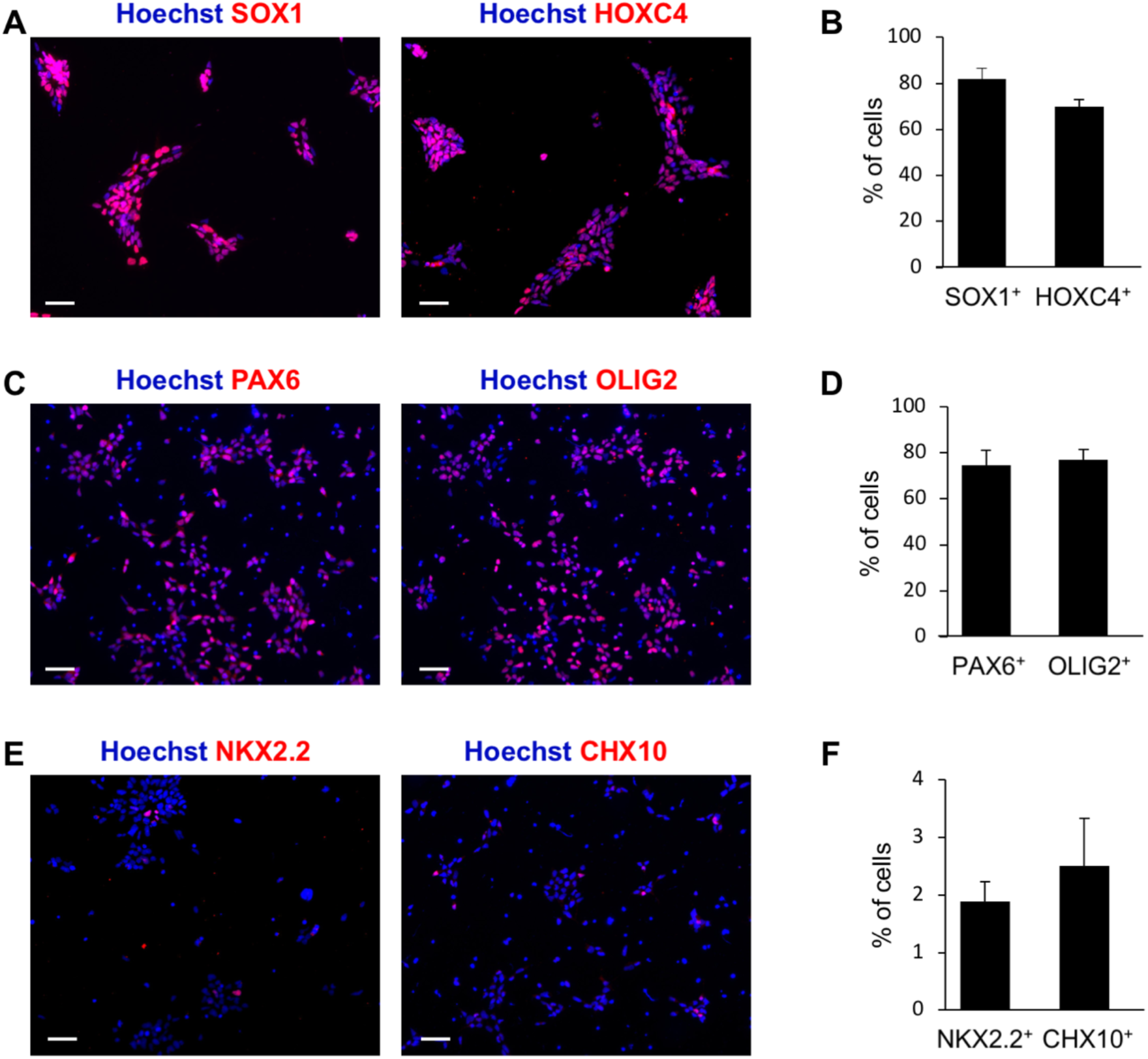
Generation of an enriched population of human iPSC-derived motor neuron progenitor cells. (A-B) Representative images (A) and quantification (B) of neural progenitor cells (NPCs) subjected to immunocytochemistry with either anti-SOX1 or anti-HOXC4 antibodies 6 days after the start of the induction protocol. (C-D) Representative images (C) and quantification (D) of MNPCs subjected to immunocytochemistry with either anti-OLIG2 or anti-PAX6 antibodies 12 days after the start of the induction protocol. (E-F) Representative images (E) and quantification (F) of V3 interneuron progenitors (INPs) subjected to immunocytochemistry with anti-NKX2.2 antibody or V2 INPs visualized with anti-CHX10 antibody, 12 days after the start of the induction protocol. For all graphs, n = 3 cultures (with >500 cells in random fields for each culture). Scale bars, 50 μm.

Continued exposure of MNPCs to RA and purmorphamine led to the generation of an enriched population of induced cells (67±7%) expressing the typical pan-MN markers ISL1 and HB9 in combination (Amoroso et al., 2013) (Figure 2A and B), as early as 25 days after the start of the differentiation protocol. Across a number of different MN derivation experiments, the yield of ISL1^+^/HB9^+^ cells varied from ∼40% to as high as ∼80% (Figure 2B), possibly the result of different MN survival rates from one culture to another, and/or variability of cell culture reagents from one lot to another. The majority (68±11%) of iPSC-derived ISL1^+^/HB9^+^ MNs did not express the LHX3 protein (Thaler et al., 2002; Agalliu et al., 2009) and instead expressed FOXP1, commonly considered as a marker of limb innervating MNs of the lateral motor column (LMC) (Dasen et al., 2003; 2008; Amoroso et al., 2013) (Figure 2C and D). Among the ISL1^+^/HB9^+^ cells that did not express FOXP1, 15±3% did not express LHX3 either, suggesting that this fraction of MNs had a molecular profile (FOXP1^-^/LHX3^-^) similar to that of MNs of the hypaxial motor column (HMC) containing MNs innervating respiratory muscles *in vivo* (data not shown). Consistent with this possibility, we observed that 10±2% of ISL1^+^/HB9^+^ MNs expressed the transcription factor POU CLASS 3 HOMEOBOX 1/OCTAMER BINDING PROTEIN 6/SCIP (SCIP), known to be enriched *in vivo* in MNs of the phrenic motor column (PMC), a component of the HMC (Bermingham et al., 1996; Philipidou et al., 2014; Machado et al., 2014) (Figure 2C and D). Lastly, only 11±4% of the h-iPSC-derived MNs were FOXP1^-^ and expressed LHX3, thus exhibiting a profile (FOXP1^-^/LHX3^+^) resembling that of MNs of the medial motor column (MMC) (Figure 2C and D). Together, these observations suggest that h-iPSC-derived MNPCs can be differentiated into a population of heterogenous spinal MNs comprising cells exhibiting molecular profiles suggestive of LMC (68%), MMC (11%), and HMC (15%) MN fates, thus highlighting the ability of h-iPSCs to give rise to heterogenous MN cultures.

**Figure 2.**
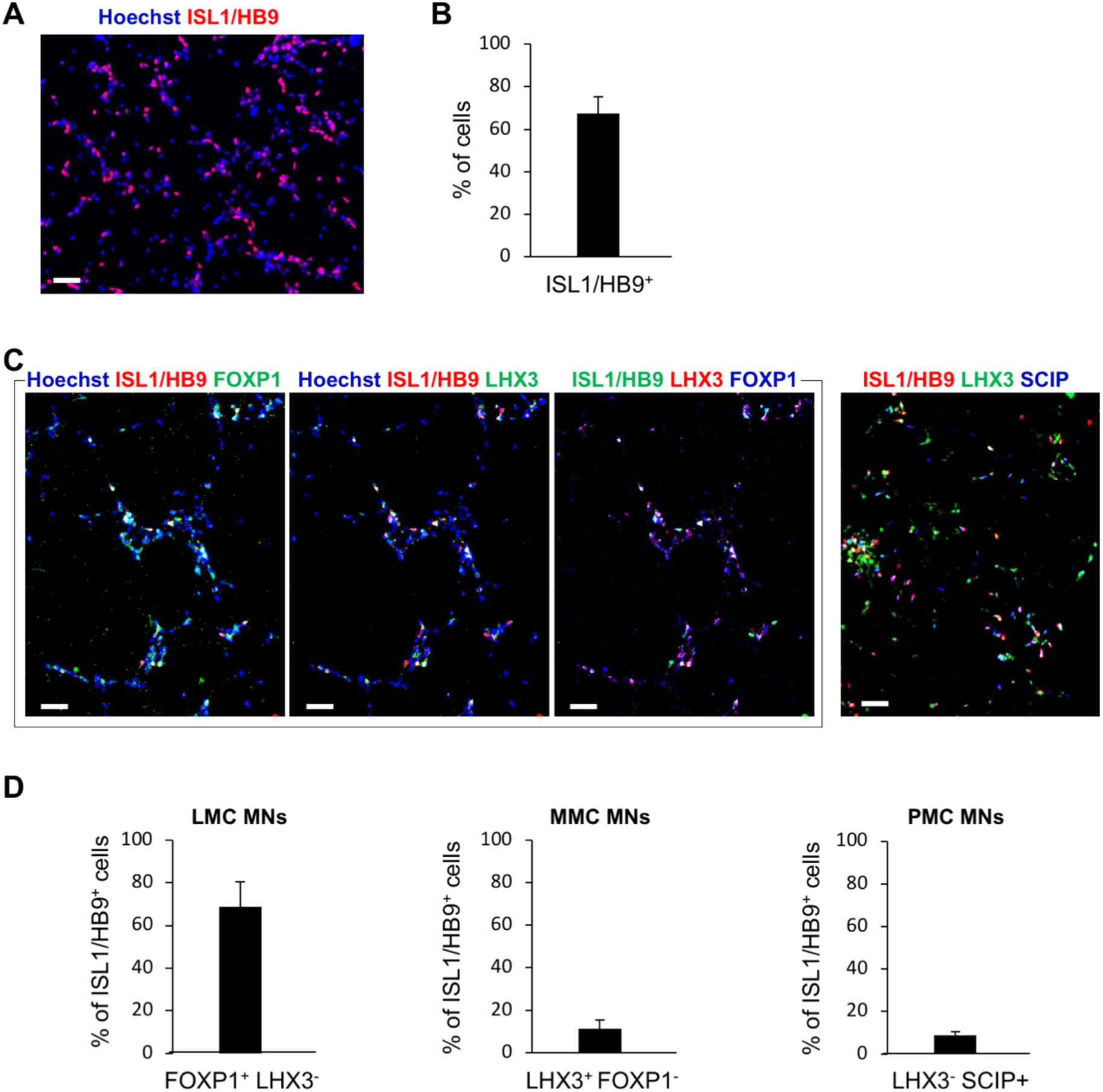
Different subpopulations of human iPSC-derived motor neurons identified by immunocytochemistry. (A-B) Representative image (A) and quantification (B) of h-IPSC-derived MNs visualized by immunocytochemistry with the pan-MN markers HB9 and ISL1 combined (ISL1/HB9) 25 days after the start of the induction protocol (n = 5 cultures; >500 cells in random fields for each culture). (C-D) Representative images (C) and quantification (D) of MN subpopulations identified through expression of different combinations of anti-ISL1/HB9, anti-LHX3, anti-FOXP1, and anti-SCIP antibodies, as revealed by immunocytochemistry. ISL1^+^/HB9^+^/FOXP1^+^/LHX3^-^ cells are defined as LMC MNs. ISL1^+^/HB9^+^/LHX3^+^/FOXP1^-^ cells are defined as MMC MNs. ISL1^+^/HB9^+^/SCIP^+^/LHX3^-^ cells are defined as PMC MNs. For all graphs in (D), n = 3 cultures (with >500 cells in random fields for each culture). LMC, lateral motor column, MMC, medial motor column, PMC, phrenic motor column. Scale bars, 50 μm.

### Characterization of cellular heterogeneity of human iPSC-derived MN culture using single-cell RNA sequencing

To further characterize the cellular composition of h-iPSC-derived MN cultures, we performed microdroplet-based sc-RNAseq, which enables a precise characterization of the genome-wide expression profile of individual cells (Stegle et al., 2015). We chose to analyze day-28 MN cultures, because at this relatively early stage of *in vitro* differentiation, induced MNs have not yet begun to coalesce into large cell clusters (as they do at later stages) that would make it technically challenging to isolate single cells. After sc-RNAseq, FASTQ files were acquired from 5,900 single cells. We used the Bioconductor software (Lun et al., 2016) to perform quality control steps to filter out damaged cells, followed by principal-component analysis and t-distributed stochastic neighbor embedding (t-SNE) analysis of the remaining cells. A total of 4,584 cells passed quality control, with a median of 5,421 mapped reads per cell and a median of 2,215 genes expressed per cell. The percentage of mitochondrial genes present in most cells was less than 2%.

The quality-controlled 4,584 cells were divided into 14 shared nearest neighbour graph-based clusters (scran) that exhibited distinct gene expression patterns, consistent with cellular heterogeneity in the h-iPSC-derived MN culture (Figure 3A). As shown in Figure 3B, the majority of induced cells were mainly contained in clusters #4 (28% of the cells), #6 (15%), #10 (10%) and #11 (9%). In order to characterize the different cell types in the identified clusters, we performed integrative analysis to compare the cell proportions and gene expression differences between each cluster. Based on differential genes enriched in each cluster, 6 major types of cells were identified (Figure 3C). Clusters #1, #3 and #11 contained a large proportion of cells identified as MNPCs or MNs, based on their expression of specific markers such as *NEUROGENIN2* (*NEUROG2*), *OLIG2, HB9, ISL1, ISL2*, and *CHOLINE ACETYL TRANSFERASE* (*CHAT)*. Cells of cluster #13 were annotated as “undifferentiated stem-like cells” with a high expression of the pluripotency markers, *OCTAMER BINDING PROTEIN 4* (*OCT4*) and *NANOG* (Nichols & Smith, 2012) (Figure 3D). Clusters #10 and #12 were annotated as “NPCs” based on their expression of *SOX1, SOX2* and *MARKER OF PROLIFERATION KI-67* (*MKI67*) (Kan et al., 2007; Graham et al., 2003; Scholzen & Gerdes, 2000). Clusters #5 and #6 were annotated as “INs” with high expression of specific marker genes for IN progenitors such as *SOX14, NKX2*.*2, VISUAL SYSTEM HOMEOBOX 1* (*VSX1*), *CHX10*, and *SINGLE-MINDED HOMOLOG 1* (*SIM1*) (Ericson et al., 1997; Novitch et al., 2001; Sugimori et al., 2007; Alaynick et al. 2011; Debrulle et al., 2019), as well as the mature IN marker *CALBINDIN 1* (*CALB1*) (Alvarez et al., 2005). Cluster #2 was annotated as “astrocytic glial cells” based on high expression of *S100 CALCIUM BINDING PROTEIN B* (*S100β*) and *SOX9* (Rosengren et al., 1986, Sun et al., 2017). Cluster #4 represented a mix of MNPCs, astrocytic glial cells and oligodendrocytes, with high expression of *NEUROG2, OLIG2, HB9, ISL1, ISL2, VESICULAR ACETYLCHOLINE TRANSPORTER* (*VACTH), S100β* and *SOX9*, as well as *PLATELET DERIVED GROWTH FACTOR RECEPTOR ALPHA* (*PDGFRα*) and *GALACTOSYLCERAMIDASE* (*GALC*) (Hall et al., 1996; Miller, 2002). Clusters #8, #9 and #14 represented “apoptotic cells”, with high expression of *BCL2 ASSOCIATED X PROTEIN* (*BAX*) (Wei et al., 2001; D’Orsi et al., 2015) and *NERVE GROWTH FACTOR* (*NGF*) (Frade and Barde, 1999).

**Figure 3:**
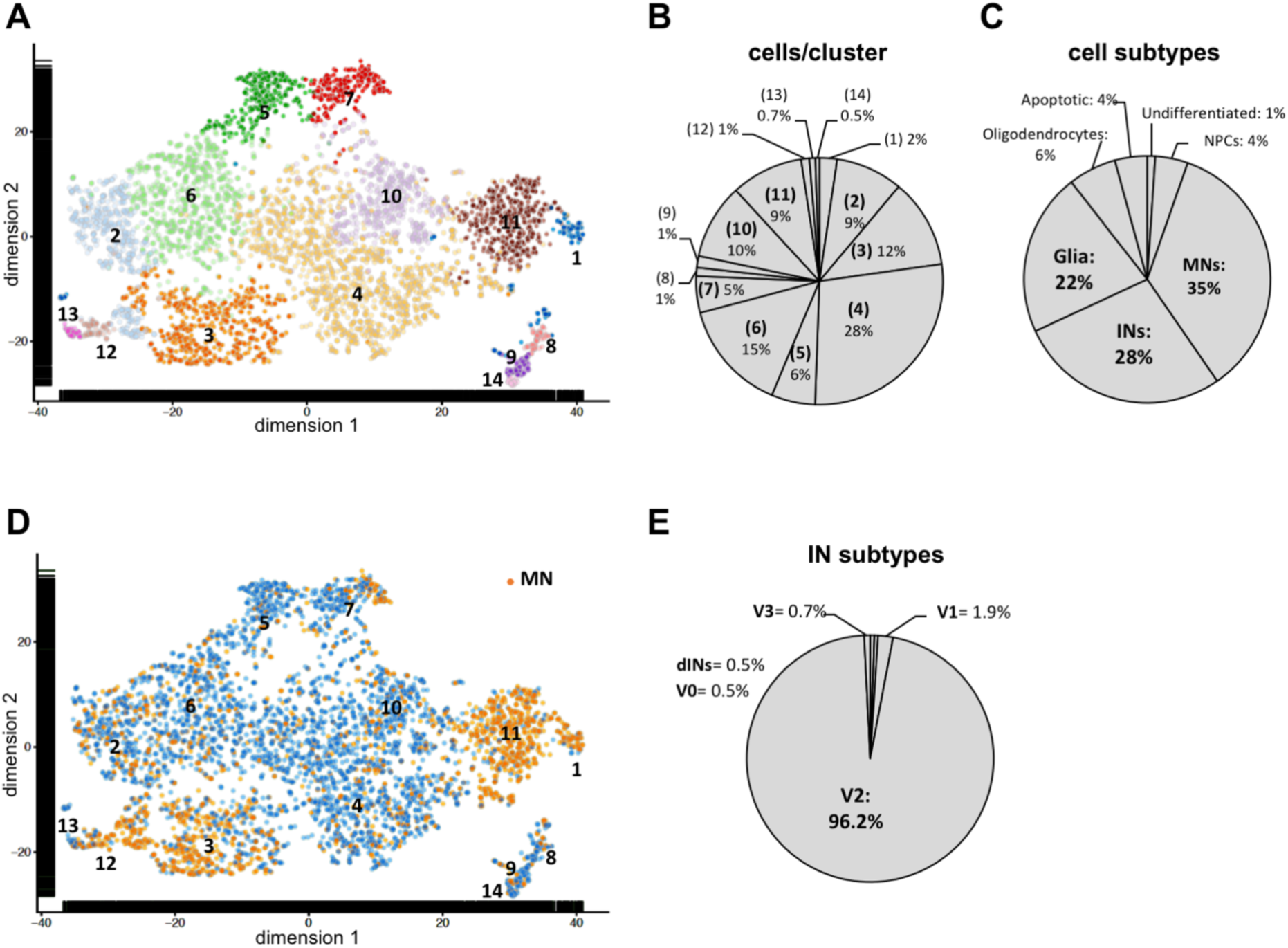
Heterogenous composition of human iPSC-derived motor neuron culture defined by single cell RNA sequencing. (A) t-SNE plot of h-iPSC derived MN culture after 28 days of differentiation, showing 14 cell shared nearest neighbour graph-based cell clusters. (B) Percentage of total cells present in each cluster. (C) Relative proportion of iPSCs, NPCs, INs, MNs and glial cells, identified based on the specific expression of the following genes: *NANOG* and *OCT4* for iPSCs; *SOX1, SOX2*, and *MKI67* for NPCs; *PAX2, PAX3, LBX1, EVX1, EN1, CHX10, GATA3, SOX14, SIM1, TLX3* for INs; *OLIG2, NEUROG2, HB9, ISL1, ISL2, CHAT* for MNs; *S100B, SOX9* for glial cells. (D) t-SNE plot colored for MNPCs and MNs (orange dots), identified based on the specific expression of the following genes: *NEUROG2, HB9, ISL1, ISL2, CHAT*. Blue dots represent the rest of the cells. (E) Proportion of each of the spinal interneuron (IN) subtypes identified based on combinatorial gene expression patterns: dorsal INs (dINs), *LBX1, PAX2, TLX3*; V0, *EVX1*^+^/*EN1*^-^; V1, *EVX1*^-^/*EN1*^+^; (V2), *CHX10, SOX14, GATA3*; (V3) *SIM1, NKX2*.*2*.

To confirm this cluster annotation, we correlated genes specifically expressed in different clusters with defined gene ontology (GO) terms. Genes that were expressed in clusters associated with NPCs were significantly enriched for neurogenesis (p = 3.366e^-07^) and nervous system development (p = 8.511e^-05^) categories. Genes that were specifically expressed in clusters associated with MNs were significantly enriched for MN axon guidance pathways (p = 3.084e^-05^). Genes that were specifically expressed in the IN clusters were significantly enriched for ventral spinal cord IN specification (p = 8.826e^-05^) and neuronal differentiation (p = 3.004e^-06^). Genes that were specifically expressed in the astrocytic glial cell clusters were significantly enriched for immune system process (p = 6.116e^-06^) and immune responses (p = 8.909e^-05^). Genes that were enriched in clusters associated with apoptotic cells were significantly enriched for regulation of programmed cell death (p = 1.039e^-10^) and neuronal apoptotic processes (p = 7.76e^-05^). Overall, the induced culture was mainly enriched in MNs (35%), INs (28%) and glial cells (22%) (Figure 3D). As shown in Figure 3E, the vast majority (96.2%) of INs were identified as V2 INs expressing *SOX14, CHX10* and *GATA BINDING PROTEIN 3* (*GATA3*). Ventral V3 INs, expressing *NKX2*.*2* and *SIM1*, represented only 0.7% of the IN population, and the more dorsal IN populations were almost absent, specifically V1 INs [*ENGRAILED1 (EN1)*^+^/*EVEN-SKIPPED HOMEOBOX 1* (*EVX1)*^-^] accounted for 1.9% of cells; V0 INs (*EN1*^-^/*EVX1*^+^) for 0.5%; and dorsal INs [*LADYBIRD HOMEOBOX 1* (*LBX1*)^+^/*T CELL LEUKEMIA HOMEOBOX 3* (*TLX3*)^+^, or *LBX1*^+^/*PAX2*^+^) for 0.5% of cells. These results show that h-iPSC-derived cultures obtained under MN differentiation experimental conditions and subjected to sc-RNAseq contain mainly MNs and V2 INs. Furthermore, when compared to immunocytochemistry results shown in Figures 1 and 2, as well as previous similar immunocytochemistry studies by other groups (e.g., Amoroso et al., 2013; Maury et al., 2014; Du et al., 2015), these findings suggest that MNs viability may be affected by the single-cell isolation procedure more than other cell types, leading to a more selective loss of MNs during sc-RNAseq compared to other cell types.

### Different subtypes of human iPSC-derived MNs identified by single-cell RNA sequencing

We next sought to identify the diverse MN subtypes derived from h-iPSCs. To this end, we selected the cells collectively characterized as MNPCs and MNs, with high expression of *NEUROG2, OLIG2, HB9, ISL1, ISL2*, and *CHAT*, as described above. A total of 1,629 such cells passed quality control, with a median of 6,322 mapped reads per cell and a median of 2,483 genes expressed per cell. The percentage of mitochondrial genes present in most cells was less than 2%. The 1,629 cells were divided into 8 shared nearest neighbour graph-based clusters exhibiting distinct gene expression patterns, suggesting heterogeneity in the MNPC/MN population (Figure 4A). As shown in Figure 4B, the majority of cells were mainly found in clusters #2 (26% of the cells), #1 (23%), #5 (20%) and #6 (16%).

**Figure 4:**
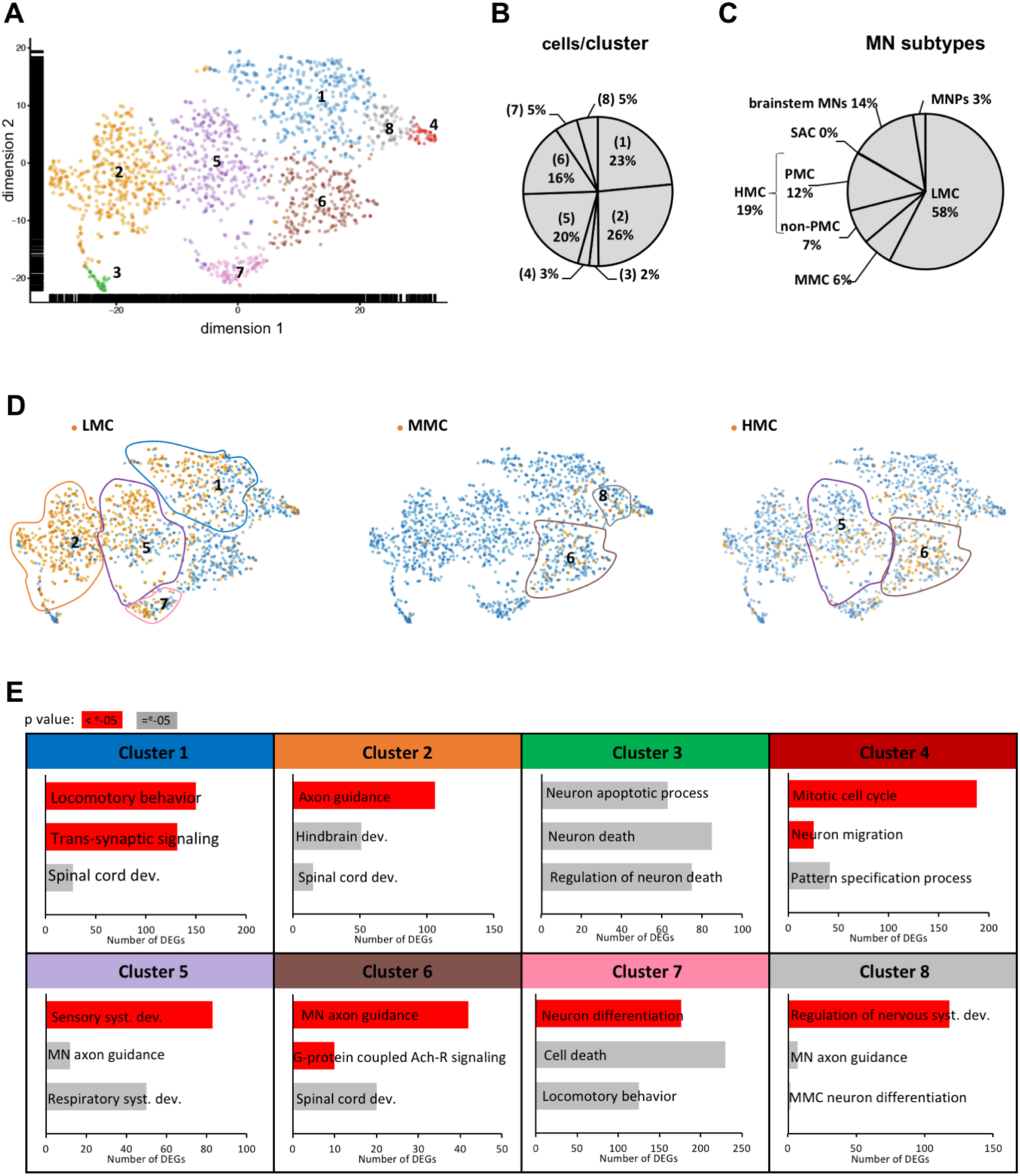
Identification of motor neuron subtypes with single-cell RNA sequencing. (A) t-SNE plot of all cells identified as MNs in h-iPSC derived MN culture after 28 days of differentiation, showing 8 shared nearest neighbour graph-based cell clusters. (B) Percentage of total cells present in each cluster. (C) Proportion of each MN subtype, identified by their expression of specific genes: *OLIG2* (MNPCs); *FOXP1*^+^/*LHX3*^-^ (LMC); *FOXP1*^-^/*LHX3*^+^ (MMC); *FOXP1*^-^/*LHX3*^-^/*HOXA5*^+^ (cervical HMC); cervical HMC expressing *SCIP* and/or *TSHZ1* (PMC); *PHOX2B*^+^ (SAC); *OTX2, MAFB* (brainstem MNs). (D) t-SNE plot colored for the 3 major subtypes of spinal MNs (orange dots): LMC (*FOXP1*^+^/*LHX3*^-^); MMC (*FOXP1*^-^/*LHX3*^+^); and cervical HMC, defined as *FOXP1*^-^/*LHX3*^-^/*HOXA5*^+^. (E) Gene Ontology (GO) analysis of the most differentially expressed genes between each cluster. Three «Biological Process» GO categories with a p value <e-05 are shown for each cluster. The number of differentially expressed genes (DEGs) associated with each category is reported, and each category is colored in grey if p = e^-05^ or red if p <e^-05^. dev., development; syst., system; Ach-R= acetylcholine receptor.

To identify the different MN subtypes within these clusters, we performed integrative analysis to compare the cell proportions and gene expression differences between each cluster. Based on differential genes enriched in each cluster, 6 major types of cells were identified (Figure 4C). Clusters #1, #2, #5 and #7 harboured a large proportion of cells identified as LMC MNs, based on their expression of *FOXP1* and lack of expression of *LHX3* (Figure 4D, top t-SNE plot). Conversely, MMC MNs expressing *LHX3* and lacking *FOXP1* expression were mainly contained in clusters #8 and #6 (Figure 4D, middle t-SNE plot). Furthermore, spinal HMC MNs expressing *HOMEOBOX A5* (*HOXA5*) and lacking both *FOXP1* and *LHX3* were found in clusters #5 and #6 (Figure 4D, bottom t-SNE plot). In addition, cluster #4 represented MNPCs with higher expression of *NKX6*.*1, OLIG2* and *LHX3* (Briscoe et al., 2000), while cluster #3 was annotated as “apoptotic cells”, with higher expression of *BAX, CASPASE 3* (*CASP3*), *NGF* and *FAS CELL SURFACE DEATH RECEPTOR* (*FAS*) (Raoul et al., 1999).

To confirm this cluster annotation, we searched for genes specifically expressed in each cell type that were enriched for the expected appropriate gene ontology (GO) terms (Figure 4E). For example, genes that were specifically expressed in the MNPC cluster were significantly associated with mitotic cell cycle (p = 2.596e^-49^), neuronal migration (p = 3.193e^-07^), and pattern specification processes (p = 7.509e^-05^). Genes that were specifically expressed in the apoptotic cell cluster were significantly enriched for neuronal apoptotic process (p = 6.596e^-05^) and regulation of neuronal death (p = 6.111e^-05^). Genes that were specifically expressed in the LMC clusters were significantly enriched for locomotor behavior (p = 9.601e^-09^), trans-synaptic signaling (p = 1.341e^-07^) and spinal cord development (p = 7.078e^-05^). Genes that were specifically expressed in the MMC clusters were significantly enriched for MN axon guidance (p = 3.192e^-07^), G-protein coupled acetylcholine receptor signaling pathways (p = 3.192e^-07^) and spinal cord development (p = 4.941e^-05^). Genes that were specifically expressed in the HMC clusters were significantly enriched for sensory system development (p = 3.835e^-06^), MN axon guidance (p = 8.383e^-05^) and respiratory system development (p = 7.284e^-05^). Interestingly, MNs expressing the specific brainstem marker genes, *ORTHODENTICLE HOMEOBOX* 2 (*OTX2*) and *MAF BZIP TRANSCRIPTION FACTOR B* (*MAFB*) were also identified and were mainly contained in cluster #2. Cluster #2 was indeed significantly enriched for hindbrain development (p = 2.307e-^05^) as well as spinal cord development (p = 5.027e^-05^), suggesting the presence of both brainstem and spinal cord MNs in this cluster. Overall, LMC MNs were the most predominant (58%), followed by HMC MNs (19%), and brainstem MNs (14%). The majority of HMC MNs (63%) expressed genes such as *SCIP* and *TEASHIRT ZINC FINGER HOMEOBOX 1* (*TSHZ1*), known to be enriched in phrenic MNs, and they were thus annotated as PMC MNs (Figure 4C). MNPCs were also detected (6%), whereas we did not detect MNs of the spinal accessory column (SAC) (Figure 4C). Together, these results provide evidence for different subtypes of h-iPSC-derived MNs, as well as other neural cell types that normally arise during the transition from undifferentiated progenitors to committed precursors to differentiated neurons in the spinal cord.

## Discussion

In most studies to date, the characterization of h-iPSC-derived MNs has relied mainly on the expression of specific molecular markers by either immunocytochemistry or real-time PCR approaches (Li et al. 2005; Patani et al. 2011; Amoroso et al. 2013; Chen et al. 2014; Maury et al. 2014; Du et al. 2015). While no single marker is completely MN-specific *in vivo*, nascent spinal MNs are characterized by transient co-expression of LIM homeodomain (LIM-HD) transcription factors like HB9, ISL1/2, and LHX3 (Sharma et al., 1998; Arber et al., 1999; Thaler et al., 1999). Thus, most MN derivation studies rely on a combination of HB9 and ISL1/2 immunostaining (referred to as pan-MN staining -Amoroso et al., 2013) to characterize the MN phenotype *in vitro*. However, LIM-HD proteins are gradually down-regulated during development *in vivo*, suggesting that these proteins may not be ideal markers for more mature MNs *in vitro*. The genes, *CHAT* and *VACTH* are expressed by all cholinergic neurons including spinal MNs and they have been used as markers for more developmentally mature MNs (Habecker & Landis, 1994). These generic MN markers are frequently used in combination with more specific motor column MN markers, such as FOXP1 or SCIP, to identify particular MN subtypes. Even these combined approaches, however, fall short of providing strategies for precisely defining the molecular profiles of all h-iPSC-derived MNs, especially when considering that common differentiation protocols give rise to heterogenous neural cell cultures (Li et al., 2005; Patani et al., 2011; Amoroso et al., 2013; Chen et al., 2014; Maury et al., 2014; Du et al., 2015). Based on these observations, the present study aimed at developing protocols to perform untargeted characterization of gene expression in all cell subtypes present in cultures of h-iPSC derived spinal MNs.

We chose to analyze day-28 MN cultures because, at this relatively early stage of *in vitro* differentiation, induced MNs already express the typical pan-MN markers ISL1 and HB9. Furthermore, at this stage of differentiation, h-iPSC-derived MNs have not yet coalesced into large cell clusters, as they typically do at more developmentally mature stages *in vitro*. The presence of these large MN clusters would make it technically challenging to isolate single MNs without causing cell damage due to the need of harsher dissociation techniques. The sc-RNAseq results presented in this study show that h-iPSCs can give rise, in 28 days of differentiation, to heterogenous neural cell cultures containing a majority of hindbrain and spinal MNs, as well as other spinal neural cells such as INs and glia. We observed that immunocytochemical analysis of h-iPSC-derived cultures after 25 days of differentiation showed a relative proportion of induced MNs (67%) larger than the fraction of MNs detected using sc-RNAseq (35%). This situation could be the result of a specific loss of MNs over other cell types during single-cells suspension preparation prior to sequencing, since mechanical and enzymatic cell dissociation may damage axons and dendrites, likely leading to increased cell death of MNs.

Strategies to generate and quantify h-iPSC-derived MN diversity are important to be able to study the function and dysfunction of different human MNs, including their pathologies in *in vitro* models of MN diseases such as ALS (Frey et al. 2000; Stifani, 2014). A number of studies have begun to characterize different types of h-iPSC-derived MNs, including limb-innervating LMC MNs and axial muscle-innervating MMC MNs, based on transcriptional profiles first characterized in the developing mouse spinal cord (Dasen et al., 2009; Alaynick et al., 2011). The *FOXP1* gene is enriched in LMC MNs (Dasen et al., 2003 and 2008; Amoroso et al., 2013), while the expression of LHX3 characterizes mainly mature MMC MNs *in vivo* (Thaler et al., 2002; Agalliu et al., 2009). While most studies (e.g., Maury et al., 2014; Davis-Dusenbery et al., 2014; Qu et al., 2014) reported mixed populations of LMC and MMC subtypes in human stem cell-derived MNs *in vitro*, Patani and colleagues showed enhanced specification of the MMC fate with the omission of retinoids from the differentiation medium (Patani et al., 2011). Conversely, Amoroso and coworkers showed an enrichment in LMC MN identity with the combined use of RA, the SHH pathway activator Smoothened Agonist, and purmorphamine (Amoroso et al., 2013). Additional MN column-specific markers are needed, as LHX3, for example, is expressed by all MNs early in development, but is maintained specifically in MMC subtypes when they mature *in vivo* (Sharma et al., 1998; Thaler et al., 2002). Furthermore, HMC subtypes remain poorly characterized in h-iPSC-derived MN cultures.

Together, the present immunocytochemistry and sc-RNAseq data strongly suggest that h-iPSCs can give rise to spinal MNs of the lateral, medial, hypaxial and phrenic motor columns *in vitro*. Both approaches reveal that the majority (58%) of MNs exhibit a molecular profile similar to that of LMC MNs, consistent with the enrichment in LMC MN identity previously obtained with a similar differentiation protocol using RA and purmorphamine (Amoroso et al., 2013). Consistent with previous reports (Maury et al., 2014), MNs with a molecular profile similar to that of brainstem MNs were also obtained from h-iPSCs in the present study. Furthermore, our data suggest that MNs exhibiting a molecular profile corresponding to HMC MNs represent 19% of the h-iPSC-derived MNs, and that 12% of these cells express either *SCIP* or *TSHZ1*, two PMC MN markers. In summary, the results of these studies show that untargeted assessment of gene expression by RNA profiling on a single cell level can provide a global measure of spinal MN diversity in h-iPSC-derived MN cultures.

Improved means to precisely phenotype the different MN subtypes generated from human iPSCs are expected to promote advancements in various fields of research. They will facilitate the study of the biology of different types of human MNs, enabling a better characterization of their physiological properties. Moreover, they may lead to the development of strategies to enrich for defined MN subtypes that may offer more attractive options for modeling specific motor neuron diseases and/or injury conditions. In turn, deeply phenotyped disease-relevant MN subtypes may provide ideal experimental model systems to develop cellular assays with increased potential to facilitate early-stage drug discovery efforts in ALS and other MN diseases.

## Acknowledgments

We thank Yeman Tang, Carol Chen, and Rita Lo for invaluable advice and assistance. These studies were funded in part by the Douglas Avrith MNI-Cambridge Neuroscience Collaboration Initiative (SS, SP, RH), the Canadian Institutes for Health Research and Fonds de la recherche en Sante-Quebec (SS), and the Tony Proudfoot Postdoctoral Fellowship Award (LT).

## Author Contributions

LT performed all experiments, data analysis, figure preparation and wrote the first draft of the manuscript. LT and RH analyzed the sc-RNAseq data. SS, TD and SP conceived the study plan. SS designed the experiments and supervised data analysis and manuscript writing.

## References

Agalliu, D., Takada, S., Agalliu, I., McMahon, A. P., & Jessell, T. M. (2009). Motor neurons with axial muscle projections specified by Wnt4/5 signaling. Neuron, 61(5), 708–720.

Alaynick, W. A., Jessell, T. M., & Pfaff, S. L. (2011). SnapShot: Spinal cord development. Cell, 146(1), 178–178.e1.

Alizadeh, A., Dyck, S. M., & Karimi-Abdolrezaee, S. (2019). Traumatic spinal cord injury: An overview of pathophysiology, models and acute injury mechanisms. Front. Neurol., 10, 282.

Alvarez, F. J., Jonas, P. C., Sapir, T., Hartley, R., Berrocal, M. C., Geiman, E. J., Goulding, M. (2005). Postnatal phenotype and localization of spinal cord V1 derived interneurons. J. Comp. Neurol., 493(2), 177–192.

Amin, N., Tan, X., Ren, Q., Zhu, N., Botchway, B. O. A., Hu, Z., & Fang, M. (2019). Recent advances of induced pluripotent stem cells application in neurodegenerative diseases. Prog. Neuropsychopharmacol. Biol. Psychiatry, 95, 109674.

Amoroso, M. W., Croft, G. F., Williams, D. J., O’Keeffe, S., Carrasco, M. A., Davis, A. R., Wichterle, H. (2013). Accelerated high-yield generation of limb-innervating motor neurons from human stem cells. J. Neurosci., 33(2), 574–586.

Arber, S., Han, B., Mendelsohn, M., Smith, M., Jessell, T. M., & Sockanathan, S. (1999). Requirement for the homeobox gene Hb9 in the consolidation of motor neuron identity. Neuron, 23(4), 659–674.

Ardhanareeswaran, K., Mariani, J., Coppola, G., Abyzov, A., & Vaccarino, F. M. (2017). Human induced pluripotent stem cells for modelling neurodevelopmental disorders. Nat. Rev. Neurol., 13(5), 265–278.

Bermingham, J. R., Jr, Scherer, S. S., O’Connell, S., Arroyo, E., Kalla, K. A., Powell, F. L., & Rosenfeld, M. G. (1996). Tst-1/Oct-6/SCIP regulates a unique step in peripheral myelination and is required for normal respiration. Genes Dev., 10(14), 1751–1762.

Briscoe, J., Pierani, A., Jessell, T. M., & Ericson, J. (2000). A homeodomain protein code specifies progenitor cell identity and neuronal fate in the ventral neural tube. Cell, 101(4), 435–445.

Burke, A. C., Nelson, C. E., Morgan, B. A., & Tabin, C. (1995). Hox genes and the evolution of vertebrate axial morphology. Development, 121(2), 333–346.

Chen, H., Qian, K., Du, Z., Cao, J., Petersen, A., Liu, H., Zhang, S. C. (2014). Modeling ALS with iPSCs reveals that mutant SOD1 misregulates neurofilament balance in motor neurons. Cell Stem Cell, 14(6), 796–809.

Dasen, J. S., De Camilli, A., Wang, B., Tucker, P. W., & Jessell, T. M. (2008). Hox repertoires for motor neuron diversity and connectivity gated by a single accessory factor, FoxP1. Cell, 134(2), 304–316.

Dasen, J. S., & Jessell, T. M. (2009). Hox networks and the origins of motor neuron diversity. Curr. Top. Dev. Biol., 88, 169–200.

Dasen, J. S., Liu, J. P., & Jessell, T. M. (2003). Motor neuron columnar fate imposed by sequential phases of hox-c activity. Nature, 425(6961), 926–933.

Davis-Dusenbery, B. N., Williams, L. A., Klim, J. R., & Eggan, K. (2014). How to make spinal motor neurons. Development, 141(3), 491–501.

Debrulle, S., Baudouin, C., Hidalgo-Figueroa, M., Pelosi, B., Francius, C., Rucchin, V., Clotman, F. (2019). Vsx1 and Chx10 paralogs sequentially secure V2 interneuron identity during spinal cord development. Cell. Mol. Life Sci. 2019 Dec 10.

D’Orsi, B., Kilbride, S. M., Chen, G., Perez Alvarez, S., Bonner, H. P., Pfeiffer, S., Prehn, J. H. (2015). Bax regulates neuronal Ca2+ homeostasis. J. Neurosci., 35(4), 1706–1722.

Du, Z. W., Chen, H., Liu, H., Lu, J., Qian, K., Huang, C. L., Zhang, S. C. (2015). Generation and expansion of highly pure motor neuron progenitors from human pluripotent stem cells. Nat. Commun., 6, 6626.

Ebert, A. D., Yu, J., Rose, F. F., Jr, Mattis, V. B., Lorson, C. L., Thomson, J. A., & Svendsen, C. N. (2009). Induced pluripotent stem cells from a spinal muscular atrophy patient. Nature, 457(7227), 277–280.

Egawa, N., Kitaoka, S., Tsukita, K., Naitoh, M., Takahashi, K., Yamamoto, T., Adachi, F., Kondo, T., Okita, K., Asaka, I., Aoi, T., Watanabe, A., Yamada, Y., Morizane, A., Takahashi, J., Ayaki, T., Ito, H., Yoshikawa, K., Yamawaki, S., Suzuki, S., Watanabe, D., Hioki, H., Kaneko, T., Makioka, K., Okamoto, K., Takuma, H., Tamaoka, A., Hasegawa, K., Nonaka, T., Hasegawa, M., Kawata, A., Yoshida, M., Nakahata, T., Takahashi, R., Marchetto, M.C., Gage, F.H., Yamanaka, S., Inoue, H. (2012). Drug screening for ALS using patient-specific induced pluripotent stem cells. Sci. Transl. Med., 4(145), 145ra104.

Ericson, J., Rashbass, P., Schedl, A., Brenner-Morton, S., Kawakami, A., van Heyningen, V., Briscoe, J. (1997). Pax6 controls progenitor cell identity and neuronal fate in response to graded shh signaling. Cell, 90(1), 169–180.

Frade, J. M., & Barde, Y. A. (1999). Genetic evidence for cell death mediated by nerve growth factor and the neurotrophin receptor p75 in the developing mouse retina and spinal cord. Development, 126(4), 683–690.

Francius, C., & Clotman, F. (2014). Generating spinal motor neuron diversity: A long quest for neuronal identity. Cell. Mol. Life Sci., 71(5), 813–829.

Frey, D., Schneider, C., Xu, L., Borg, J., Spooren, W., & Caroni, P. (2000). Early and selective loss of neuromuscular synapse subtypes with low sprouting competence in motoneuron diseases. J. Neurosci., 20(7), 2534–2542.

Graham, V., Khudyakov, J., Ellis, P., & Pevny, L. (2003). SOX2 functions to maintain neural progenitor identity. Neuron, 39(5), 749–765.

Habecker, B. A., & Landis, S. C. (1994). Noradrenergic regulation of cholinergic differentiation. Science (New York, N.Y.), 264(5165), 1602–1604.

Hall, A., Giese, N. A., & Richardson, W. D. (1996). Spinal cord oligodendrocytes develop from ventrally derived progenitor cells that express PDGF alpha-receptors. Development, 122(12), 4085–4094.

Haston, K. M., & Finkbeiner, S. (2016). Clinical trials in a dish: The potential of pluripotent stem cells to develop therapies for neurodegenerative diseases. Ann. Rev. Pharmacol. Toxicol., 56, 489–510.

Jessell, T. M. (2000). Neuronal specification in the spinal cord: Inductive signals and transcriptional codes. Nat. Rev. Genet., 1(1), 20–29.

Kan, L., Jalali, A., Zhao, L. R., Zhou, X., McGuire, T., Kazanis, I., Kessler, J. A. (2007). Dual function of Sox1 in telencephalic progenitor cells. Dev. Biol., 310(1), 85–98.

Kanning, K. C., Kaplan, A., & Henderson, C. E. (2010). Motor neuron diversity in development and disease. Ann. Rev. Neurosci., 33, 409–440.

Kiskinis, E., Sandoe, J., Williams, L. A., Boulting, G. L., Moccia, R., Wainger, B. J., Eggan, K. (2014). Pathways disrupted in human ALS motor neurons identified through genetic correction of mutant SOD1. Cell Stem Cell, 14(6), 781–795.

Li, X. J., Du, Z. W., Zarnowska, E. D., Pankratz, M., Hansen, L. O., Pearce, R. A., & Zhang, S. C. (2005). Specification of motoneurons from human embryonic stem cells. Nat. Biotechnol., 23(2), 215–221.

Lun, A. T., McCarthy, D. J., & Marioni, J. C. (2016). A step-by-step workflow for low-level analysis of single-cell RNA-seq data with bioconductor. F1000research, 5, 2122.

Machado, C. B., Kanning, K. C., Kreis, P., Stevenson, D., Crossley, M., Nowak, M., Lieberam, I. (2014). Reconstruction of phrenic neuron identity in embryonic stem cell-derived motor neurons. Development, 141(4), 784–794.

Maury, Y., Come, J., Piskorowski, R. A., Salah-Mohellibi, N., Chevaleyre, V., Peschanski, M., Nedelec, S. (2015). Combinatorial analysis of developmental cues efficiently converts human pluripotent stem cells into multiple neuronal subtypes. Nat. Biotechnol., 33(1), 89–96.

McCarthy, D.J., Campbell, K.R., Lun, A.T.L., Willis, Q.F. (2017). “Scater: pre-processing, quality control, normalisation and visualisation of single-cell RNA-seq data in R.” Bioinformatics, 33, 1179–1186.

Methot, L., Soubannier, V., Hermann, R., Campos, E., Li, S., & Stifani, S. (2018). Nuclear factor-kappaB regulates multiple steps of gliogenesis in the developing murine cerebral cortex. Glia, 66(12), 2659–2672.

Miller, R. H. (2002). Regulation of oligodendrocyte development in the vertebrate CNS. Prog. Neurobiol., 67(6), 451–467.

Nichols, J., & Smith, A. (2012). Pluripotency in the embryo and in culture. Cold Spring Harb. Perspect. Biol., 4(8), a008128.

Nijssen, J., Comley, L. H., & Hedlund, E. (2017). Motor neuron vulnerability and resistance in amyotrophic lateral sclerosis. Acta Neuropathol., 133(6), 863–885.

Novitch, B. G., Chen, A. I., & Jessell, T. M. (2001). Coordinate regulation of motor neuron subtype identity and pan-neuronal properties by the bHLH repressor Olig2. Neuron, 31(5), 773–789.

Patani, R., Hollins, A. J., Wishart, T. M., Puddifoot, C. A., Alvarez, S., de Lera, A. R., Chandran, S. (2011). Retinoid-independent motor neurogenesis from human embryonic stem cells reveals a medial columnar ground state. Nat. Commun., 2, 214.

Philippidou, P., Walsh, C. M., Aubin, J., Jeannotte, L., & Dasen, J. S. (2012). Sustained Hox5 gene activity is required for respiratory motor neuron development. Nat. Neurosci., 15(12), 1636–1644.

Prasad, A., & Hollyday, M. (1991). Development and migration of avian sympathetic preganglionic neurons. J. Comp. Neurol., 307(2), 237–258.

Qu, Q., Li, D., Louis, K. R., Li, X., Yang, H., Sun, Q., Wang, F. (2014). High-efficiency motor neuron differentiation from human pluripotent stem cells and the function of islet-1. Nat. Commun., 5, 3449.

Raoul, C., Henderson, C. E., & Pettmann, B. (1999). Programmed cell death of embryonic motoneurons triggered through the fas death receptor. J. Cell Biol., 147(5), 1049–1062.

Rosengren, L. E., Kjellstrand, P., Aurell, A., & Haglid, K. G. (1986). Irreversible effects of dichloromethane on the brain after long term exposure: A quantitative study of DNA and the glial cell marker proteins S-100 and GFA. Br. J. Ind. Med., 43(5), 291–299.

Scholzen, T., & Gerdes, J. (2000). The ki-67 protein: From the known and the unknown. J. Cell. Physiol., 182(3), 311–322.

Stegle, O., Teichmann, S. A., & Marioni, J. C. (2015). Computational and analytical challenges in single-cell transcriptomics. Nat. Rev. Genet., 16(3), 133–145.

Stifani, N. (2014). Motor neurons and the generation of spinal motor neuron diversity. Front. Cell. Neurosci., 8, 293.

Sugimori, M., Nagao, M., Bertrand, N., Parras, C. M., Guillemot, F., & Nakafuku, M. (2007). Combinatorial actions of patterning and HLH transcription factors in the spatiotemporal control of neurogenesis and gliogenesis in the developing spinal cord. Development, 134(8), 1617–1629.

Sun, W., Cornwell, A., Li, J., Peng, S., Osorio, M. J., Aalling, N., Nedergaard, M. (2017). SOX9 is an astrocyte-specific nuclear marker in the adult brain outside the neurogenic regions. J. Neurosci., 37(17), 4493–4507.

Thaler, J., Harrison, K., Sharma, K., Lettieri, K., Kehrl, J., & Pfaff, S. L. (1999). Active suppression of interneuron programs within developing motor neurons revealed by analysis of homeodomain factor HB9. Neuron, 23(4), 675–687.

Thaler, J. P., Lee, S. K., Jurata, L. W., Gill, G. N., & Pfaff, S. L. (2002). LIM factor Lhx3 contributes to the specification of motor neuron and interneuron identity through cell-type-specific protein-protein interactions. Cell, 110(2), 237–249.

Tsuchida, T., Ensini, M., Morton, S. B., Baldassare, M., Edlund, T., Jessell, T. M., & Pfaff, S. L. (1994). Topographic organization of embryonic motor neurons defined by expression of LIM homeobox genes. Cell, 79(6), 957–970.

Vallstedt, A., Muhr, J., Pattyn, A., Pierani, A., Mendelsohn, M., Sander, M., Ericson, J. (2001). Different levels of repressor activity assign redundant and specific roles to Nkx6 genes in motor neuron and interneuron specification. Neuron, 31(5), 743–755.

Wei, M. C., Zong, W. X., Cheng, E. H., Lindsten, T., Panoutsakopoulou, V., Ross, A. J., Korsmeyer, S. J. (2001). Proapoptotic BAX and BAK: A requisite gateway to mitochondrial dysfunction and death. Science, 292(5517), 727–730.

Wichterle, H., Lieberam, I., Porter, J. A., & Jessell, T. M. (2002). Directed differentiation of embryonic stem cells into motor neurons. Cell, 110(3), 385–397.

